# AimSeg: a machine-learning-aided tool for axon, inner tongue and myelin segmentation

**DOI:** 10.1101/2023.01.02.522533

**Authors:** Ana Maria Rondelli, Jose Manuel Morante-Redolat, Peter Bankhead, Bertrand Vernay, Anna Williams, Pau Carrillo-Barberà

## Abstract

Electron microscopy (EM) images of axons and their ensheathing myelin from both the central and peripheral nervous system are used for assessing myelin formation, degeneration (demyelination) and regeneration (remyelination). The g-ratio is the gold standard measure of assessing myelin thickness and quality, and traditionally is determined from measurements done manually from EM images – a time-consuming endeavour with limited reproducibility. These measurements have also historically neglected the innermost uncompacted myelin sheath, known as the inner myelin tongue. Nonetheless, the inner tongue has been shown to be important for myelin growth and some studies have reported that certain conditions can elicit its enlargement. Ignoring this fact may bias the standard g-ratio analysis, whereas quantifying the uncompacted myelin has the potential to provide novel insights in the myelin field. In this regard, we have developed AimSeg, a bioimage analysis tool for axon, inner tongue and myelin segmentation. Aided by machine learning classifiers trained on tissue undergoing remyelination, AimSeg can be used either as an automated workflow or as a user-assisted segmentation tool. Validation results show good performance segmenting all three fibre components, with the assisted segmentation showing the potential for further improvement with minimal user intervention. This results in a considerable reduction in time for analysis compared with manual annotation. AimSeg could also be used to build larger, high quality ground truth datasets to train novel deep learning models. Implemented in Fiji, AimSeg can use machine learning classifiers trained in ilastik. This, combined with a user-friendly interface and the ability to quantify uncompacted myelin, makes AimSeg a unique tool to assess myelin growth.

**Author Summary:** Myelin is formed by specialised cells that wrap themselves around axons and has a major role in the function, protection, and maintenance of nerves. These functions are disturbed by demyelinating diseases, such as multiple sclerosis. In this work we present AimSeg, a new tool based on artificial intelligence algorithms (machine learning) to assess myelin growth on electron microscopy images. Whereas standard metrics and previous computational methods focus on quantifying compact myelin, AimSeg also quantifies the inner myelin tongue (uncompacted myelin). This structure has been largely overlooked despite the fact that it has an important role in the process of myelin growth (both during development and in the adult brain) and recent studies have reported morphological changes associated with some diseases. We report the performance of AimSeg, both as a fully automated approach and in an assisted segmentation workflow that enables the user to curate the results on-the-fly while reducing human intervention to the minimum. Therefore, AimSeg stands as a novel bioimage analysis tool that meets the challenges of assessing myelin growth by supporting both standard metrics for myelin evaluation and the quantification of the uncompacted myelin in different conditions.

## Introduction

The myelin sheath allows faster, saltatory conduction of nerve impulses along the underlying axon without the need to increase axon diameter [1,2]. Moreover, this lipid-rich insulating layer also provides structural protection and metabolic support to the underlying axons [3]. The myelin sheath consists of plasma membrane from either oligodendrocytes (in the central nervous system, CNS) or Schwann cells (in the peripheral nervous system, PNS) wrapped around axons, and is discontinuous around their length, separated by nodes of Ranvier [4]. These cells extend cytoplasmic-filled membrane processes that are guided to reach and ensheath the axon. Myelin growth occurs by the wrapping of the leading edge of the myelin membrane process (henceforth the inner tongue) around the axon, progressing underneath the previously deposited membrane in concert with the lateral extension of the individual myelin layers along the axons [5]. Myelin compaction is initiated after a few wraps, occurring first in the outermost myelin layer and progressively spreading inwards, lagging behind the inner tongue to avoid its premature compaction. During developmental myelination, the inner tongue is enlarged but it narrows as myelin matures [4,6]. Once active myelination is completed, a smaller inner tongue remains in adult myelinated fibres [5,7], except as recently discovered in the context of some diseases [8].

Generally, most myelinated fibres have a ratio of axon to fibre diameters (g-ratio) close to the optimal value for conduction velocity of neural electrical impulses, estimated from theoretical models in the PNS and the CNS [9,10]. Additionally, larger diameter axons have more myelin wraps (thicker myelin sheath) and a lower g-ratio [11–13]. The g-ratio is widely utilised by the scientific community as a functional and structural index of optimal axonal myelination, and for assessing remyelination following myelin loss. Defects in myelination in the CNS can be assessed in this way in neurodevelopmental disorders [14–17], demyelinating diseases (e.g., multiple sclerosis, MS) [18,19], neurodegenerative diseases [20–22], as well as in rodent models of myelin abnormalities [8,23–26]. Moreover, the remyelination process in MS is characterised by thinner myelin sheaths for the diameter of the axon, giving higher g-ratios [27,28].

G-ratios are commonly calculated on EM images of chemically fixed samples, though progress has been made to try and measure these *in vivo* on MR brain scans in humans [19,29]. Despite the wide applicability and functional relevance of this metric, it neglects the inner tongue. In fact, the inner tongue in adult myelinated fibres has been largely overlooked, as it can only be detected with the high resolution provided by EM. Furthermore, these structures tend to shrink and collapse in chemically fixed and dehydrated tissue [29]. Importantly, an enlarged inner tongue will bias the standard g-ratio analysis by overestimating the diameter of the axon, thus researchers are adopting alternative ways to perform the g-ratio analysis to assess myelination/remyelination. For example, a corrected g-ratio accounting for the enlarged inner tongue has been recently proposed [26]; however, no bioimage analysis methods are available for such approaches. Moreover, advances in sample preparation for EM provide a better preservation of myelin ultrastructure [29], enabling researchers to better study the uncompacted myelin and leading to novel discoveries in myelin biology. For example, recent studies have reported an enlarged or abnormal inner tongue in transgenic mice (e.g. 2′,3′-cyclic nucleotide 3′-phosphodiesterase (CNP)-deficient mice [23,30], in conditional knock-out of activin co-receptor *Acvr1b* (31)[31] and of *Pten* (5)[5] in oligodendrocytes), and in animal models of autoantibody-mediated- and cuprizone-induced-demyelinating disease [8,25], suggested to be secondary to stressed axons with a compensatory increase in need for metabolic support from the oligodendrocyte via the inner tongue.

Several bioimage analysis approaches have been developed to analyse myelin thickness [32–38]. Many of these approaches are often implemented in semi-automated workflows that frequently require several post-processing steps. Deep learning approaches have also been applied [38,39] to segment the individual fibres and their corresponding compacted myelin. Additionally, there are methods available for the analysis of 3D EM images [40]. However, their wide application by researchers has been limited, as still only few are publicly available, well documented, and/or the code made accessible through open-source licensing. Therefore, analysis of myelin thickness from EM images is still largely performed manually by investigators, which is time-consuming and prone to selection bias, thus contributing to limited reproducibility. Notably, all the above-mentioned methods ignore the inner tongue and do not support its quantification.

This has motivated us to develop an open-access tool, named AimSeg, for the <u>seg</u>mentation of the <u>a</u>xon, the inner tongue, and the compact <u>m</u>yelin from EM data. Our goal has been to enable a more thorough assessment of the myelin sheath thickness, while decreasing the need for manual annotation, and saving time. AimSeg uses supervised machine learning methods implemented in ilastik [41] to improve the segmentation of the fibre components, and combines automated image processing with interactive user-editing stages in Fiji [42]. AimSeg automatically stores all the generated regions of interest (ROIs) in different subsets of axonal components interrelated between them and the results table by the axon IDs. The workflow code for AimSeg – initially written in the form of an ImageJ macro, and subsequently reimplemented as a Groovy script for improved performance and maintainability – is open-source and the pre-trained ilastik classifiers are publicly available together with user documentation.

## Results

### AimSeg workflow

Unlike other approaches that enable the analysis of images acquired throughout a wide range of imaging techniques [43], our goal was to develop a method to separate and analyse each of the fibre components (the axon, the compact myelin and the inner tongue) from EM images (see Fig 1). To this end, it is necessary to outline the axon, the innermost and the outermost compact myelin layers. AimSeg achieves this through the segmentation of three objects with a hierarchical relationship: the fibre cross-section (including the compact myelin sheath), the inner compact myelin layer (ICML) that consists of the axon and the inner tongue, and the axon (without the inner tongue). The combination of these masks allows the calculation of both the standard g-ratio and the inner tongue area. Our strategy relies on supervised machine learning methods, which have been demonstrated to be useful to analyse complex images such as those acquired through EM [44,45]. AimSeg can be applied as a fully automated image processing workflow or enable an assisted segmentation approach that includes interactive user-editing. Our workflow makes use of open-source bioimage analysis software (ilastik [41] and Fiji [42]). The AimSeg core pipeline is a Fiji script that takes as an input a series of files previously generated using machine learning classifiers trained using ilastik (see Fig 2).

**Fig 1.**
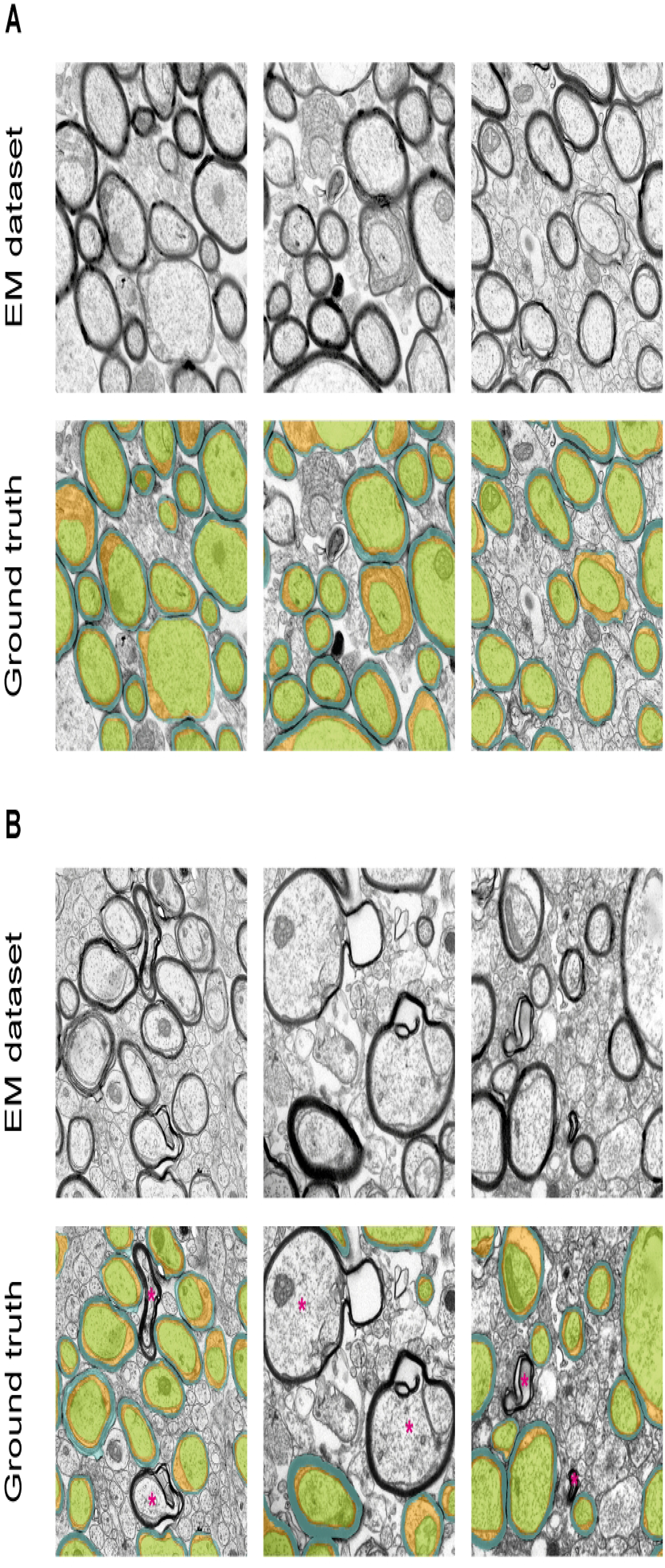
Expert-annotated ground truth for the segmentation of fibre cross-sections on electron microscopy images separating the myelin sheath components (compact myelin and inner tongue) from each other and from the axon. Examples of the dataset, which comprises transmission electron microscopy (TEM) images of the corpus callosum from adult mice **(A, top)**. The ground truth was annotated by an expert by drawing three region of interest (ROI) sets. These represent both the instance segmentation of the fibre cross-sections and the semantic segmentation of the compacted myelin (cyan), the innermost uncompacted myelin that faces the axon (also known as the inner cytoplasmic tongue, orange), and the axon (green) **(A, bottom)**. Conventional TEM sample preparation may damage the ultrastructure of some fibres, thus producing artefacts that are not of interest for quantification. Accordingly, when detected, such structures were not included within the ground truth **(B)**. Magenta asterisks indicate fibre cross-sections whose ultrastructure has been altered by the chemical fixation process.

**Fig 2.**
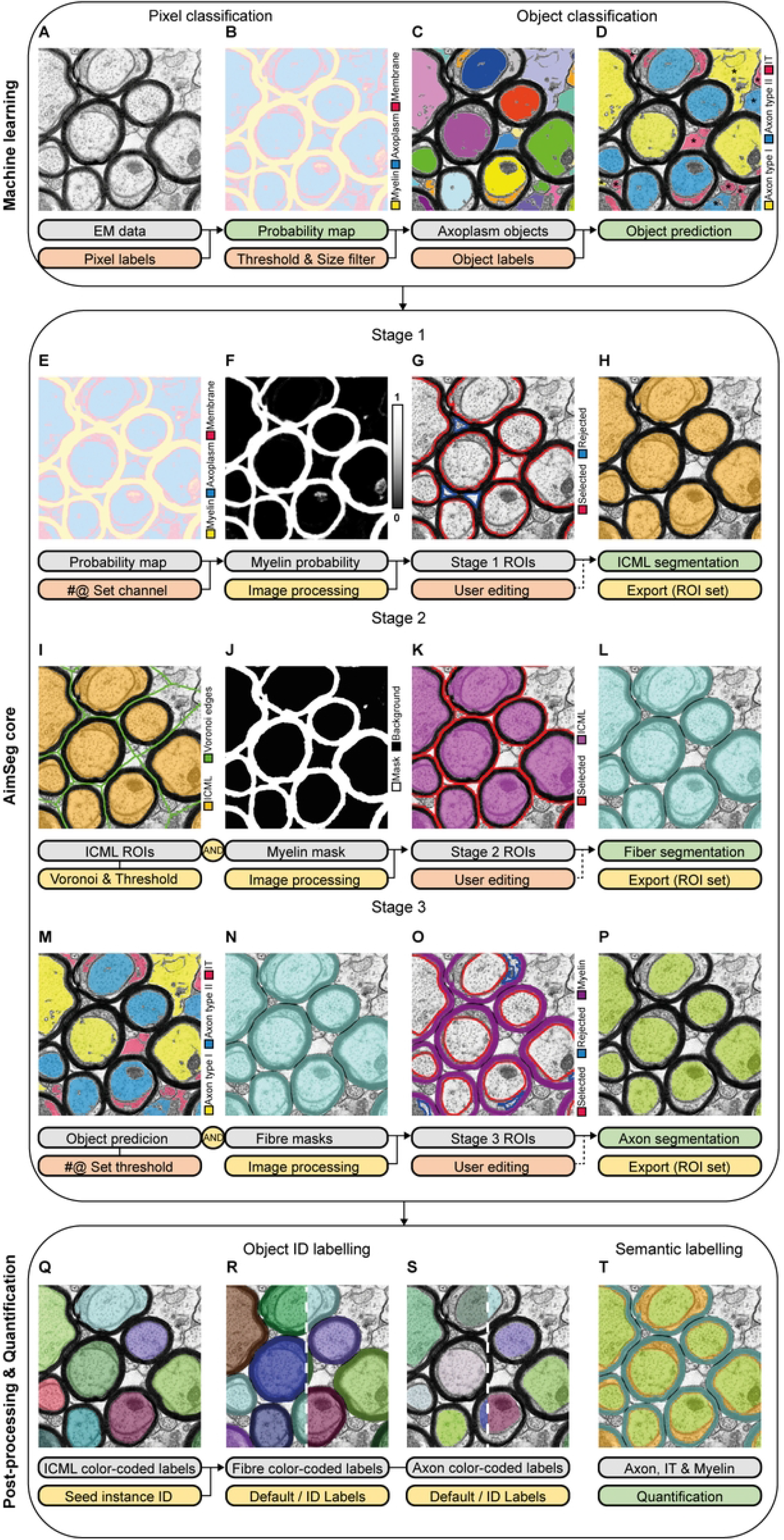
Main steps of the AimSeg bioimage analysis workflow including the optional, human-supervised steps for assisted segmentation. The AimSeg workflow is divided into three main pipelines: machine learning (ML) **(A-D)**, AimSeg core **(E-P)**, and post-processing and quantification **(Q-T)**. AimSeg requires the combination of two ML classifiers (for pixel and object classification respectively) that can be trained using, e.g., ilastik. First, the pixel classifier uses the electron microscopy (EM) data **(A)** to generate a probability map **(B)** in which each pixel is assigned a score according to its probability of belonging to three different classes: myelin, axoplasm or membrane. Then, a threshold is applied to segment the axoplasm probabilities and the smaller objects are filtered out **(C)**. The object classifier scores each object as an axon or inner tongue (IT) **(D)**. In this example two different axon classes (bigger: type I, smaller: type II) were defined, though the user can decide to include as many classes as they deem necessary for their data. Note that those objects outside myelinated fibre cross-sections (marked with an asterisk) will not affect the analysis, since they will be eliminated. The AimSeg core follows the ML steps **(E-P)**. An assisted-segmentation approach combines automated processing with steps that allow the user to amend the regions of interest (ROIs) proposed by AimSeg (user editing marked by an asterisk). Conversely, the automated mode just skips the user editing steps. The AimSeg core, either human-assisted or automated, is subdivided in three stages, each of which segments an element of the myelinated fibres; the combination of these elements will allow the quantification of the myelin, the IT and the axon. Stage 1 **(E-H)** uses the ML probability map **(E)** to get the myelin probability channel **(F)**, which is processed to perform the segmentation of the inner compact myelin layer (ICML), i.e., the axon plus the IT (namely, the fibre without the compact myelin). At this point, AimSeg presents two different types of ROIs: selected or rejected. Note that the ROI selection at Stage 1 includes a segmentation error (false positive), which is marked with an asterisk **(G)**. The user can easily toggle the ROI class (selected/rejected) or use the selection tools to add/edit ROIs. The definitive ICML-ROI set **(H)** is stored after the user-editing stage. Stage 2 **(I-L)** uses the ICML-ROIs to calculate their corresponding voronoi territories **(I)**. These are used to split the myelin binary mask **(J)** into independent, ring-shaped objects corresponding to the compact myelin. The rings are filled and post-processed to get the fibre ROIs **(K)**. Then AimSeg waits for the user to edit the selection before the definitive fibre-ROI set **(L)** is stored. Stage 3 **(M-P)** uses the ML object prediction to create two subsets of binary masks, thresholding either the objects labelled as axons or those considered to belong to other classes (IT in this example) **(M)**. The fibre ROIs **(N)** are used to ensure that only those axons wrapped by myelin (i.e., overlapping a fibre ROI) are selected. As in Stage 1, AimSeg generates two types of axon ROIs, selected, or rejected, that the user can toggle **(O)**. Once the user is satisfied with the selection, the axon-ROI set **(P)** is stored. Finally, AimSeg includes a post-processing pipeline **(Q-T)** that is crucial for data quantification, as it establishes a meaningful relationship between the three ROIs that belong to the same fibre cross-section (i.e., ICML, fibre and axon). Each ICML ROI is assigned a 3-digit code and is used as a seed **(Q)** to detect their corresponding – though unidentified – fibre **(R, left)** and axon **(S, left)** ROIs. Then, fibre and axon ROIs are labelled after the ICML code **(R-S, right)**. Once all the ROIs are labelled, quantification is performed using the semantic segmentation of each fibre cross-section **(T)**. Node colour-code: yellow (image processing), green (export), orange (user input). Note that the user intervention during the ML pipeline is only needed for training the classifiers. #@ denotes Fiji initialisation parameters.

The workflow starts with two classifiers (pixel and object classification; see Fig 2A-D) trained using supervised machine learning methods implemented within ilastik – although AimSeg has the potential to use probability maps and object predictions generated by means of any other machine learning toolkit. For this work, we used two ilastik workflows for pixel and object classification trained on transmission electron microscopy (TEM) images acquired on remyelinating tissue; specifically, the dataset includes images of the corpus callosum (CC) of adult mice after a demyelinating lesion was induced. Users can directly apply the ready-to-use classifiers pre-trained for this work, improve them by adding their own raw data and annotations, or train new classifiers from scratch by following the guidelines provided within the AimSeg documentation. We kept the pixel and object classification as independent pipelines to save RAM and speed up the process.

The assisted segmentation pipeline (see Fig 2E-P), implemented within Fiji, asks the user to select an EM file to be analysed. AimSeg requires the machine learning output to be stored in the working directory or to provide a pre-trained classifier. The pipeline is divided in three sequential stages aimed to segment each of the three fibre components from each of the fibre cross-sections in the image. In Stage 1, the myelin probability map is processed to segment the ICML (see Fig 2E-H), which is used as a seed to split the myelin sheath of neighbouring axons and filled to get the whole fibres in Stage 2 (see Fig 2I-L). Finally, axons are segmented from the machine learning object predictions in Stage 3 (see Fig 2M-P).

Additionally, after the automated steps, each stage includes an optional user-editing step that allows manual amendment of any segmentation inaccuracies before proceeding to the next stage. At these points, the user can delete ROIs, use the Fiji selection tools to edit or add new ROIs or use a series of shortcuts provided in AimSeg to interact with the ROIs while reducing the user intervention. Additionally, those ROIs filtered out during the automated processing are also shown as rejected-mode ROIs at Stage 1 and 3, so the user can toggle them to the selected-mode without drawing them from scratch (see Fig 2G and 2O).

Once the three ROI sets have been generated, an automated pipeline within the Fiji-implemented workflow is set to both post-process the three ROI sets and to extract the quantitative features (see Fig 2Q-T). The post-processing step aims to: i) remove any residual pixels that may have been left by the user during the partial erasing of ROIs during the manual editing step; ii) ensure that, as expected, the ROIs of the innermost elements do not overflow into the outer ones (e.g., the axon ROI should never break through the ICML ROI); iii) duplicate the ICML-ROI in case an axon ROI has not been selected for a fibre because of a thinner inner tongue, removing the need for the user to copy it manually; and iv) assign an ID to each of the ROIs in order to bind the different components belonging to the same fibre between the three ROI sets (see Fig 2Q-S). The latter step is important for performing a meaningful quantification, and enables the user to trace results from the final measurements table back to the image dataset. Finally, the area of each ROI of the fibre cross-section is extracted and summarised in a results table.

Users can reduce their manual intervention by annotating only one ROI set (the ICML-ROI at Stage 1) of those axons that are not suitable for morphometric quantification (e.g., because they are touching the image edge or affected by artefacts due to the EM chemical fixation process during sample preparation), as the post-processing tool decides that these fibres should not be carried forward for analysis. Therefore, these ROIs will be stored and given an ID in the results table, but with 0 values in the three columns. Thus, the row count can be used to determine the total number of fibres per image without the need to further annotate incomplete or problematic fibres in Stage 2 and 3 for which g-ratio analysis cannot be done.

### Assessment of the automated and supervised segmentation

To quantitatively assess the performance of our strategy, we compared the segmentation output obtained using our workflow with a ground truth manually annotated by an expert (see Fig 1). This was performed on TEM images of remyelinating tissue different from those used to train the classifiers. As discussed above, one critical point in our design is the user-supervised step intended to correct or edit the automatic segmentation. Therefore, to evaluate the impact of such manual steps, we compared the expert’s ground truth with two different workflow outputs: a fully automated workflow without any user intervention (see Fig 3A-B), and a supervised (assisted segmentation) workflow that included a limited version of all the user-editing steps. The supervised workflow allows the user to edit the ROI set by including or discarding ROIs automatically suggested by our tool, but not manually drawing them (see Fig 3C). This constraint was imposed to avoid the overcorrected segmentation often derived from the use of Fiji’s selection tools, and allowed us to assess the potential of AimSeg to facilitate the user-assisted segmentation.

**Fig 3.**
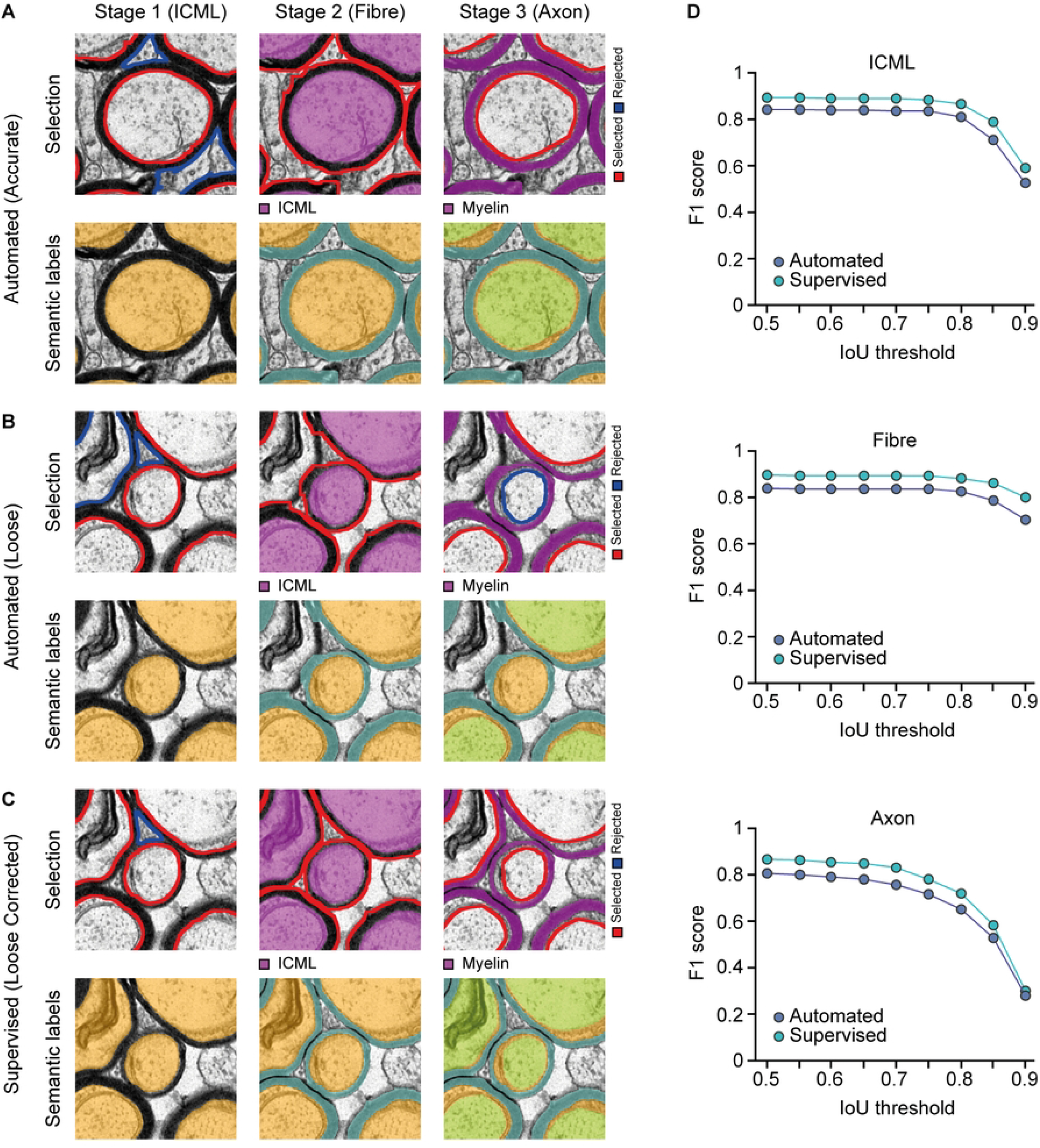
AimSeg segmentation performance, assessed independently for the three exported region of interest (ROI) sets: the inner compact myelin layer (ICML; i.e., the axon plus the inner tongue), the fibre and the axon. Evaluation of the instance segmentation performed either in a fully automated **(A, B)** or an assisted segmentation approach **(C)**. At first, since no user intervention was allowed, results included both accurate **(A)** and loose **(B)** segmentations (marked with a white asterisk). Note that skipping an ICML instance at Stage 1 caused the myelin mask of the surrounding fibres to overflow at Stage 2. Additionally, an axon instance is missing at Stage 3. For the supervised case **(C)**, the user was allowed to curate the AimSeg selection during the user-editing stages, but only by toggling the selected/rejected ROIs or using other AimSeg shortcuts to quickly modify them, but never by drawing new ROIs or manually editing the suggested ones. Note that toggling the ICML ROI rejected at Stage 1 solves the overflowing issue at Stage 2. Moreover, the axon ROI rejected at Stage 3 can be easily switched on. **(D)** Quantitation of the segmentation performance is based on the F1 score, an object-based metric, which is plotted for increasing intersection over union (IoU) thresholds for the estimation of shape matching accuracy in both the automated and the human-supervised results.

We used the ‘F1 score’ to assess the performance of AimSeg during the automated and supervised workflows. The average F1 score of all the annotated images was calculated across a range of intersections over union (IoU) thresholds, starting from 0.5 up to 0.9 (in increments of 0.05). The representation of the F1 score along an IoU threshold range allows one to simultaneously look at the correctly identified objects and the pixel-wise closeness of their corresponding ROIs. A higher F1 score denotes a good detection of the object while a lower score corresponds to a poor object detection.

F1 scores were independently computed for the fibre, the ICML, and the axon components (see Fig 3D). The automated approach (*Automated* and *Axon Autocomplete* modes on) demonstrated a considerable capability to predict the fibre constituents, with the F1 score being consistently high across the IoU thresholds. We also demonstrate that these results can be substantially improved by allowing the user to review and amend the segmented objects for all three components, even when not allowed to use the selection tools in Fiji to upgrade the ROI selection or to create new ones. For example, setting a 0.5 IoU we observe that the usage of the AimSeg shortcuts lead to similar performance improvements at the three stages: the axon F1 score increases from 0.81 to 0.87; the ICML F1 score from 0.84 to 0.89; the fibre F1 sore from 0.84 to 0.9. Indeed, looking at the F1 score improvement for the whole range of IoU thresholds computed for the assessment, we observe a modest but consistent increment for the three components. The performance of the original AimSeg macro segmentation was also assessed, with similar results (S1 Fig).

### Computation time

To assess the computation time employed with AimSeg versus the manual annotation, we first compared the time required to manually annotate one image with the time spent to process the image via the AimSeg pipeline without user-supervision (i.e., in a fully automated way). The total time required to manually annotate one image was 51.5 min; the time to automatically process one image was 19.2 s. We further assessed the computational time required for each automated step of the AimSeg core workflow. The time required for the automated processing steps (i.e., excluding parameterisation, data import and user supervision) of each stage per image was: 2.85 s (Stage 1), 9.68 s (Stage 2), and 3.52 s (Stage 3). The elapsed time for the post-processing and quantification pipeline was 3.14 s. Therefore, the computational time for automated processing across the three stages was negligible when compared with the highly time-consuming endeavour of annotating the images manually. The Groovy implementation improves the macro performance in terms of computation time, especially during the post-processing pipeline (S2 Fig).

## Discussion

The g-ratio is the gold standard for the assessment of the optimal myelination of axons. However, the calculation of this highly used metric neglects the existence of the uncompacted myelin of the inner tongue; a fibre component with relevance during myelination and remyelination, and whose thickness variation may contribute to the identification of both physiological and pathological processes. The lack of bioimage analysis tools accounting for the inner tongue makes its quantification a tedious task, requiring the manual annotation of EM images by experts: a time-consuming process prone to bias. Here we present AimSeg, a bioimage analysis tool for <u>a</u>xon, inner tongue, and <u>m</u>yelin sheath <u>seg</u>mentation of fibre-cross sections from EM images.

The AimSeg workflow uses open-source bioimage analysis software to combine supervised machine learning with an assisted segmentation pipeline to facilitate the annotation of these three fibre components. Afterwards, a post-processing pipeline amends some common annotation errors and integrates the different ROI sets before quantifying the metrics of interest. The evaluation of segmentation accuracy, assessed using the F1 score, showed AimSeg’s good performance when used in full-automated mode, which can be further improved with minimal user effort by using pre-defined shortcuts in editing mode. By employing an assisted segmentation approach, poorly segmented objects can be edited, benefiting the segmentation of subsequent stages. Furthermore, our workflow allows researchers to considerably reduce the time required for image annotation, thus facilitating the quantification of larger datasets.

Among the three fibre components, the axon was the most difficult element to segment. This is to be expected, as the axon is the last processed object within the three-step semi-automated pipeline, and its segmentation is burdened by any errors accumulated during the first two stages used to determine what defines a fibre. Additionally, axon segmentation faces other challenges such as the axolemma thinness and the presence of electron-dense bodies characteristic for mitochondria or neurofilaments within the axoplasm. Despite these limitations, AimSeg was still able to segment most axons. Nonetheless, we have identified a few factors that may affect the performance of AimSeg. Namely, some fibres that were very small (0.2–0.4 μm), non-convex, or sparsely myelinated were often overlooked or poorly segmented. We also observed a difference in the segmentation performance between well preserved fibres and those affected by tissue-processing artefacts; this highlights the importance of sample preparation that represents a common obstacle in bioimage analysis.

EM is a unique source of imaging data that can unravel the complex architecture of cells and its biological implications. However, its nanometric resolution is a double-edged sword: bioimage analysis methods are difficult to implement and their use is not yet widespread among researchers. Consequently, manual annotation can often result in partial analysis of image data. Conversely, a common bioimage analysis approach is focusing on the segmentation of specific structures or organelles, so a meaningful analysis to solve a specific biological question requires building a tailored workflow combining different bioimage analysis tools [46]. The use of machine learning and deep learning methods is common among these tools. The AimSeg workflow uses ilastik-built-in machine learning algorithms to facilitate the complex process of EM segmentation. To this aim, it takes advantage of the machine learning tools provided by ilastik (standing out for being interactive and providing a user-friendly graphical user interface (GUI) for users without coding experience), the versatility of Fiji, and the interoperability of both toolkits.

The end goal of AimSeg is to enable a more sophisticated analysis in the field of myelin biology, and to provide the researchers with bioimage analysis tools to facilitate these types of analyses. AimSeg serves both as a computer-assisted quantitative tool and an annotation aid. In fact, the need for reliably annotated datasets has significantly increased with the expansion of deep learning methods, yet highly dependent on the existence of human-annotated ground truths. Accordingly, we consider AimSeg as a powerful tool to train novel deep learning models. Indeed, amending segmentation results obtained automatically has proven to be an efficient approach to improve pre-existing models [47].

With this work, we aim to contribute to filling the gap between myelin biology and bioimage analysis. We believe that AimSeg may facilitate the study of the long-neglected inner tongue by providing a user-friendly, open-source platform for its quantification. Moreover, our assisted segmentation approach enhances the throughput capability of the analysis while enabling manual annotation. Overall, AimSeg’s features and novel metrics have the potential to support more sensitive and high-throughput approaches to analyse myelin ultrastructure beyond the standard g-ratio.

## Methods

### Dataset

Experimental protocols involving mice were performed under UK Home Office project licence PADF15B79 (A.W.) issued under the Animals (Scientific Procedures) Act. Adult mouse CC tissue was obtained and processed for EM as described in [18,48]. TEM images used in this study were collected on a JEOL JEM-1400 Plus TEM with GATAN One View camera at 7.1 K magnification with image dimensions 8.62 μm x 8.62 μm.

### Training the classifiers

Our workflow begins by training two classifiers trained using supervised machine learning tools implemented within ilastik 1.3.3post3 [41], the first for pixel classification and the second for object classification. Both classifiers were trained interactively using a subset of 4 images randomly selected as the training set.

Three different classes were defined for the pixel classification: i) compacted myelin, ii) electron-lucent cytoplasm-like regions (e.g., the axoplasm of axons, the non-compact myelin of the inner tongue or other regions corresponding to cell bodies) and iii) cell membranes, such as the axolemma or the inner tongue membrane. An expert annotated the EM dataset to provide the classifier with examples. Features were automatically selected using the Wrapper Method implemented within ilastik with a 0.5 Size Set Penalty. Output images were exported as probability maps, and together with the EM raw images constitute the input of the subsequent object classification. The classifier generates a pixel probability map for each pre-defined class.

The object classification pipeline within ilastik starts performing an instance segmentation, taking as input the cytoplasm probability map that was smoothed (0.6 sigma), thresholded (0.5 threshold) and size-filtered (with objects smaller than 2500 pixels rejected). We defined three different classes: two representing axoplasm cross-sections (larger or smaller), and the other representing inner tongue cross-sections. The rest of the objects obtained through the instance segmentation, including cells or unmyelinated axons, are not annotated during the training process, and thus their predictions are ignored. We included all the shape and intensity distribution features implemented within ilastik except for the object classification; conversely, the location features were ignored. Once trained, predicted objects are exported as labelled images.

### Assisted segmentation pipeline

The segmentation of the fibre components is performed in Fiji [42] over a pipeline that receives as input both the raw EM images and the output of the machine learning classifiers – one multi-channel file for the probability map and an image file for the object predictions. The aim is to obtain a ROI set of the fibre cross-sections plus two additional ROI sets: the ICML and the axon. Each ROI set is generated in three different stages: Stage 1 computes the ICML-ROI set, which determines the seed of the fibre, as it is expanded to include the compact myelin and return the fibre-ROI set (Stage 2). These two ROI sets enable the calculation of the standard g-ratio, which will be biased when the inner tongue is enlarged. To provide metrics for the inner tongue, however, it is necessary to get the axon-ROI set (Stage 3). Every stage begins with the automated processing of the images and provides a segmentation output (ROI sets). The AimSeg user-assisted segmentation enables investigators to curate the ROI sets on-the-fly. Stages 1 and 3 automatically suggest amendments for the ICML that the user can incorporate to speed up the user-editing process and enhance the subsequent step.

Stage 1 uses the probability map (generated with the machine learning workflow for pixel classification), specifically the channel corresponding to the compacted myelin. Myelin is thresholded to generate a binary mask, which is inverted to obtain the potential ICML instances. Initially, objects at the edge of the image are excluded and the remaining objects are smoothed by a closing operation. Then, the objects on the edges are recovered in a different binary mask. The objective is to apply different criteria to filter the objects depending on whether they are complete or not (touching the edges). Objects are filtered out depending on circularity – being more permissive with incomplete objects – and size. Selected objects are converted into ROIs to enable the user to interact with them. Moreover, rejected objects are also displayed as ROIs, as in some cases these may represent false negatives that should be included. Selected and rejected ROIs are set to belong to different groups and displayed in different colours, so the user can easily identify them and switch between the selected/rejected mode.

Stage 2 uses the ICML ROIs as a seed to calculate the Voronoi territories for each fibre. These territories are used to split the compact myelin mask and obtained at Stage 1 by segmenting the myelin probability, thus obtaining individualised, ring-shape objects. These hollow objects are filled to generate the fibre masks, and those touching the image edges are excluded; whereas the ICML objects on the edges were necessary to obtain a proper segmentation of the fibre objects, the fibre objects on the edges are no longer necessary, as they are not suitable for myelin quantification. The remaining objects are post-processed. First, the fibres are smoothed by a closing operation. Then convex objects are split using Fiji’s binary watershed. Then a binary reconstruction is performed using the ICML objects as seeds, thus recovering only the split fibre objects that partially overlap any seed. In case a fibre was split incorrectly, the corresponding objects are merged again by a closing operation. Selected objects are converted into ROIs for the user editing stage. Since fibre ROIs are not filtered by area or any other property, no rejected ROIs are suggested as potential false negatives at Stage 2.

Stage 3 uses the object prediction (generated with the machine learning workflow for object classification), selecting only those objects labelled as axons while filtering out those labelled as inner tongue (or any other class that the user may include). The selected objects are smoothed by a closing operation and filled. Then, those objects that do not overlap with any detected fibre ROI are also rejected. The selected objects are converted to ROIs and then transformed into their corresponding convex hulls. Those convex hulls overflooding the ICML ROIs are corrected. Additionally, the rejected inner tongue objects are also selected and smoothed, and those overlapping with a detected fibre ROI are displayed as rejected ROIs, similarly to what was formerly described for Stage 1.

Basic binary operations (erode, dilate, open, close) are implemented as modifications of ImageJ’s source code. Voronoi and watershed transforms are called by running the corresponding GUI commands. Binary reconstruction is part of the morphological operations provided at Fiji’s Morphology update site [49].

### Post-processing and quantification pipeline

After the assisted segmentation pipeline, a post-processing pipeline performs an automated correction and integration of the generated ROI sets. User-editing stages may lead to the introduction of artefacts within the different ROI sets. One of the most common errors is the annotation of individual or small groups of pixels isolated from the target object. Typically, this occurs as a result of partially erasing a ROI that ends up as a ShapeRoi (i.e., a composite selection). To correct this, the pipeline splits the ShapeRoi into PolygonRois (i.e., simple selections) and returns as the true ROI only the bigger one. Moreover, in order to avoid unnecessary work, AimSeg includes the *Autocomplete Axon* mode. When activated, the lack of the axon-ROI for a certain fibre cross-section is interpreted by the post-processing pipeline as the absence of inner tongue on the fibre (due to an almost total shrinkage), preventing the user from having to try to generate two identical ROIs for the ICML and the axon. Instead, an axon-ROI is automatically generated as a copy of the ICML-ROI, thus completing the partial axon-ROI set, and circumventing any conflict regarding the ICML-axon area coverage. The pipeline also assigns a fibre-specific ID to each ROI in order to establish a hierarchy between the different components belonging to the same fibre. This process is based on the overlap between the different ROI sets. Additionally, AimSeg ensures that the smaller objects are always contained by the bigger components – i.e., the ICML should never overflow the fibre, and the axon must not spill over the ICLM. This is achieved by calculating the intersection of the different ROIs labelled with the same ID. Once corrected, the area of each ROI is quantified and stored in a results table. Each row is labelled with the fibre-specific ID and the table contains three additional columns for the area values: axon, ICML and fibre.

### Expert annotations

Manual annotations of the three ROIs (axon, ICLM and fibre) were done by a single expert using the freehand selection tools within Fiji [42], thus generating the ground truth for the respective ROI set of entire images. Five randomly selected TEM images were used for the expert annotations. Each ROI set was exported as an independent ZIP archive file.

The AimSeg workflow was designed to only extract quantitative measurements from complete fibre cross-sections (i.e., fibre cross-sections touching the edges of the image are not quantified). Additionally, the optimal segmentation of all the fibre elements may be limited when conventional EM sample preparation with chemical fixation is used. This method could affect the ultrastructure of fibres, by causing the deformation of the fibre or altering its structure to an extent that precludes the recognition of its components [29]. To make the user annotations comparable with the AimSeg workflow output, both the ground truth annotations and the AimSeg output were submitted to an additional filtering using a Groovy script that deletes the ROIs corresponding to those fibre cross-sections touching the edges of the image or missing any of their three ROI sets (ICLM, fibre, or axon).

The decision not to include those fibres on the segmentation assessment was based on the fact that those fibres are not suitable for quantification and can be easily removed using a simple script.

Finally, since the proposed segmentation method is set to avoid the generation of overlapping fibre ROIs, the annotations of neighbouring fibres were kept apart from each other by means of a background wall thick enough to distinguish the independent objects in an 8-connected setting.

### Metrics for the assessment of instance segmentation

To evaluate the performance of the segmentation we used the F1 score. We computed an object-based metric rather than a pixel-based metric because the goal of our pipeline is to perform the instance segmentation of each individual fibre and its components to extract morphometric features. Briefly, the F1 score is based on computing the overlapping degree between the target (T), i.e., the ground truth annotated by an expert, and the prediction (P) masks, automatically generated by our method. Then, a specific overlap value is set to determine the threshold between correctly segmented and ill-segmented objects. The F1 score is calculated from this per-object assessment with possible values from 0 to 1. A higher F1 score corresponds to a better performance, so a value of 1 indicates the correct detection of all the objects.

First, the overlap between T and P is calculated for each object as the intersection over union (IoU) metric (also known as Jaccard index).

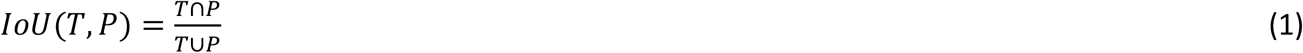

Where the intersection (T⋂P) is the count of the pixels shared by both T and P, whereas the union (T⋃P) is the count of the pxiels that are part of either T, P or both. Therefore, the IoU has a value of 1 for identical objects, while a value of 0 indicates that there is no overlap between T and P.

Then, an IoU threshold is set to label each object as a true positive (TP) if a T mask has a corresponding P mask with an overlapping degree greater than the established threshold or a false negative (FN) if a T object has no corresponding prediction mask, i.e., the overlapping degree is below the threshold; a false positive (FP) is obtained when a P mask does not match any T object according to the established IoU threshold.

Finally, the F1 score is defined as the harmonic mean of precision and recall:

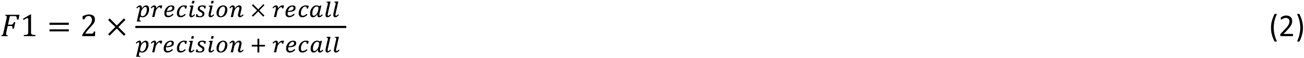

Precision is determined by the proportion of predicted objects that had a match with the annotated ground truth and is defined as:

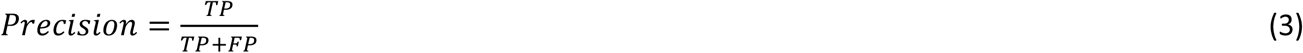

Recall is determined by the proportion of target objects that had a match on the prediction mask, calculated as:

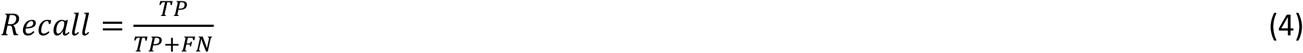

Therefore, the F1 score is calculated as:

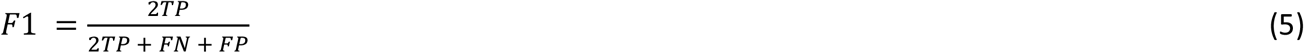

It is common to compute the F1 score along a range of IoU values [50], since the selection of a single IoU threshold may be considered an arbitrary measure. We excluded IoU values below 0.5 to avoid the conflict of pairing a T object with two P objects, or vice versa. Notably, a F1 score of 1 is practically unattainable, even when comparing the annotations of two human operators. Therefore, we used a range from 0.5 to 0.9 with increments of 0.05. This approach enables the assessment of the segmentation performance by plotting a score based on the proportion of correctly identified objects (F1) over the pixel-wise accuracy (IoU).

The precision, recall and F1 score of the model have been computed using a customised Fiji macro (https://github.com/paucabar/assess_segmentation). The script uses region connection calculus [51] to match overlapping objects between binary images. The AimSeg performance was assessed on the output generated by the latest implementation of AimSeg as a Groovy script.

### Hardware

Computation time quantification was performed on a HP OMEN 15-DC0000NS laptop with an Intel® Core™ i7-8750H processor, 16 GB of RAM and an NVIDIA® GeForce® GTX 1060 graphic card.

## Data availability

Fiji pipelines can be easily installed by means of a dedicated Fiji update site. Requirements, installation, usage documentation and scripts (including both the original macro and updated Groovy implementations) are provided in GitHub (https://github.com/paucabar/AimSeg). Additionally, pre-trained ilastik classifiers for both pixel and object classification are also available in GitHub. The validation dataset, including both TEM images and their corresponding label images are accessible in Zenodo (10.5281/zenodo.6327209).

## Acknowledgements

The authors would like to thank Stephen Mitchell for helping with the acquisition of TEM data, and Ana Domingo-Muelas and Chiara Sgattoni for helping with designing the graphics.

## Author contributions

A.M.R and P.C.B. wrote the manuscript. P.C.B. and J.M.M.R. designed the figures. A.M.R., P.C.B., A.W. and J.M.M.R. conceived the project; P.C.B. and B.V. designed the image processing strategy for the AimSeg workflow. P.C.B. developed the scripts in collaboration with P.B. A.M.R. performed the demyelination experiments, processed the samples, and acquired the TEM data. A.M.R. generated the ground truth and recorded the time for manual annotation. P.C.B implemented the pipeline to assess the segmentation performances. J.M.M.R., B.V. and A.W. supervised the project and provided expert guidance. P.C.B. wrote the AimSeg documentation. P.B., J.M.M.R. and B.V. proofread the documentation and tested AimSeg. All authors reviewed and approved the final manuscript.

## Supporting information captions

**S1 Fig. Segmentation performance of the AimSeg implementation as an ImageJ macro for the three ROI sets: (A)** the inner compact myelin layer (ICML; i.e., the axon plus the inner tongue), **(B)** the fibre and **(C)** the axon sets. The assessment was performed calculating the object-based metric known as F1 score, which is plotted for increasing intersection over union (IoU) thresholds for the estimation of shape-matching accuracy in both the automated and the human-supervised results.

**S2 Fig. Computation time of the AimSeg implementations as a Groovy script or as an ImageJ macro**. The elapsed time was computed for each of the main stages that makes up the AimSeg workflow running the Automated mode (with no stops for user supervision). We can distinguish four blocks: stage 1 (S1), segmentation of inner compact myelin layer (ICML; i.e., the axon plus the inner tongue); stage 2 (S2), segmentation of the fibre; stage 3 (S3), segmentation of the axon; post-processing and quantification (PPQ).

## References

1. Ritchie JM. On the relation between fibre diameter and conduction velocity in myelinated nerve fibres. Proc R Soc London - Biol Sci. 1982;217(1206):29–35.

2. Waxman SG. Determinants of conduction velocity in myelinated nerve fibers. Muscle Nerve. 1980;3(2):141–50.

3. Simons M, Nave KA. Oligodendrocytes: Myelination and axonal support. Cold Spring Harb Perspect Biol. 2016;8(1). doi: 10.1101/cshperspect.a020479.

4. Snaidero N, Simons M. Myelination at a glance. J Cell Sci. 2014;127(14):2999–3004.

5. Snaidero N, Möbius W, Czopka T, Hekking LHP, Mathisen C, Verkleij D, et al. Myelin membrane wrapping of CNS axons by PI(3,4,5)P3-dependent polarized growth at the inner tongue. Cell. 2014;156(1–2):277–90.

6. Michalski J-P, Kothary R. Oligodendrocytes in a Nutshell. Front Cell Neurosci. 2015;9:340. doi:10.3389/fncel.2015.00340.

7. Chang KJ, Redmond SA, Chan JR. Remodeling myelination: Implications for mechanisms of neural plasticity. Nat Neurosci. 2016;19(2):190–7.

8. Johnson ES, Ludwin SK. Evidence for a “dying-back” gliopathy in demyelinating disease. Ann Neurol. 1981;9(3):301–5.

9. Rushton WAH. A theory of the effects of fibre size in medullated nerve. J Physiol.1951;115(1):101–22.

10. Chomiak T, Hu B. What is the optimal value of the g-ratio for myelinated fibers in the rat CNS? A theoretical approach. PLoS One. 2009;4(11):e7754.doi: 10.1371/journal.pone.0007754.

11. Donaldson HH, Hoke GW. On the areas of the axis cylinder and medullary sheath as seen in cross sections of the spinal nerves of vertebrates. J Comp Neurol Psychol. 1905;15(1):1–16.

12. Franklin RJM, Ffrench-Constant C. Regenerating CNS myelin - From mechanisms to experimental medicines. Nature Reviews Neuroscience. 2017;18(12):753–69.

13. Hildebrand C, Hahn R. Relation between myelin sheath thickness and axon size in spinal cord white matter of some vertebrate species. J Neurol Sci. 1978;38(3):421–34.

14. Fields RD. White matter in learning, cognition and psychiatric disorders. Trends Neurosci. 2008;31(7):361–70.

15. Vanes LD, Moutoussis M, Ziegler G, Goodyer IM, Fonagy P, Jones PB, et al. White matter tract myelin maturation and its association with general psychopathology in adolescence and early adulthood. Hum Brain Mapp. 2020;41(3):827–39.

16. Owen JP, Marco EJ, Desai S, Fourie E, Harris J, Hill SS, et al. Abnormal white matter microstructure in children with sensory processing disorders. NeuroImage Clin. 2013;2(1):844–53.

17. Nave KA, Ehrenreich H. Myelination and oligodendrocyte functions in psychiatric diseases. JAMA Psychiatry. 2014;71(5):582–4.

18. Rittchen S, Boyd A, Burns A, Park J, Fahmy TM, Metcalfe S, et al. Myelin repair invivo is increased by targeting oligodendrocyte precursor cells with nanoparticles encapsulating leukaemia inhibitory factor (LIF). Biomaterials. 2015; 56:78–85.

19. York EN, Martin S-J, Meijboom R, Thrippleton MJ, Bastin ME, Carter E, et al. MRI-derived g-ratio and lesion severity in newly diagnosed multiple sclerosis. Brain Commun. 2021;3(4):1–10.

20. Pettit LD, Bastin ME, Smith C, Bak TH, Gillingwater TH, Abrahams S. Executive deficits, not processing speed relates to abnormalities in distinct prefrontal tracts in amyotrophic lateral sclerosis. 2013;3290–304.

21. Defrancesco M, Egger K, Marksteiner J, Esterhammer R, Hinterhuber H. Changes in White Matter Integrity before Conversion from Mild Cognitive Impairment to Alzheimer ‘s Disease. PLoS One. 2014;9(8):e106062.doi: 10.1371/journal.pone.0106062.

22. Kim HJ, Joon S, Sung H, Gon C, Kim N, Han S, et al. Alterations of mean diffusivity in brain white matter and deep gray matter in Parkinson ‘ s disease. Neurosci Lett. 2013;550:64–8.

23. Edgar JM, McLaughlin M, Werner HB, McCulloch MC, Barrie JA, Brown A, et al. Early ultrastructural defects of axons and axon-glia junctions in mice lacking expression of Cnp1. Glia. 2009;57(16):1815–24.

24. Snaidero N, Velte C, Myllykoski M, Raasakka A, Ignatev A, Werner HB, et al. Antagonistic Functions of MBP and CNP Establish Cytosolic Channels in CNS Myelin. Cell Rep. 2017;18(2):314–23.

25. Weil MT, Möbius W, Winkler A, Ruhwedel T, Wrzos C, Romanelli E, et al. Loss of Myelin Basic Protein Function Triggers Myelin Breakdown in Models of Demyelinating Diseases. Cell Rep. 2016;16(2):314–22.

26. Meschkat M, Steyer AM, Weil MT, Kusch K, Jahn O, Piepkorn L, et al. White matter integrity in mice requires continuous myelin synthesis at the inner tongue. Nat Commun. 2022;13(1).

27. Périer O, Grégoire A. Electron microscopic features of multiple sclerosis lesions. Brain. 1965;88(5):937–52.

28. Stikov N, Campbell JSW, Stroh T, Lavelée M, Frey S, Novek J, et al. In vivo histology of the myelin g-ratio with magnetic resonance imaging. Neuroimage. 2015;118:397–405.

29. Weil MT, Ruhwedel T, Meschkat M, Sadowski B, Möbius W. Transmission electron microscopy of oligodendrocytes and myelin. In: Methods in Molecular Biology. Humana Press Inc.; 2019. p. 343–75.

30. Lappe-Siefke C, Goebbels S, Gravel M, Nicksch E, Lee J, Braun PE, et al. Disruption of Cnp1 uncouples oligodendroglial functions in axonal support and myelination. Nat Genet. 2003;33(3):366–74.

31. Dillenburg A, Ireland G, Holloway RK, Davies CL, Evans FL, Swire M, et al. Activin receptors regulate the oligodendrocyte lineage in health and disease. Acta Neuropathol.2018;135(6):887–906.

32. Romero E, Cuisenaire O, Denef JF, Delbeke J, Macq B, Veraart C. Automatic morphometry of nerve histological sections. J Neurosci Methods. 2000;97(2):111–22.

33. Wang YY, Sun YN, Lin CCK, Ju MS. Segmentation of nerve fibers using multi-level gradient watershed and fuzzy systems. Artif Intell Med. 2012;54(3):189–200.

34. Liu T, Seyedhosseini M, Ellisman M, Tasdizen T. Watershed merge forest classification for electron microscopy image stack segmentation. In: Proceedings of the 21st International Conference on Pattern Recognition (ICPR2012). 11–15 Nov, 2012. Tsukaba; p. 133–7.

35. More HL, Chen J, Gibson E, Donelan JM, Beg MF. A semi-automated method for identifying and measuring myelinated nerve fibers in scanning electron microscope images. J Neurosci Methods. 2011;201(1):149–58.

36. Zhao X, Pan Z, Wu J, Zhou G, Zeng Y. Automatic identification and morphometry of optic nerve fibers in electron microscopy images. Comput Med Imaging Graph. 2010;34(3):179–84.

37. Bégin S, Dupont-Therrien O, Bélanger E, Daradich A, Laffray S, De Koninck Y, et al. Automated method for the segmentation and morphometry of nerve fibers in large-scale CARS images of spinal cord tissue. Biomed Opt Express. 2014;5(12):4145–61.

38. Zaimi A, Duval T, Gasecka A, Côté D, Stikov N, Cohen-Adad J. AxonSeg: Open source software for axon and myelin segmentation and morphometric analysis. Front Neuroinform. 2016;10:1–13.

39. Moiseev D, Hu B, Li J. Morphometric analysis of peripheral myelinated nerve fibers through deep learning. J Peripher Nerv Syst. 2019;24(1):87–93.

40. Abdollahzadeh A, Belevich I, Jokitalo E, Tohka J, Sierra A. Automated 3D Axonal Morphometry of White Matter. Sci Rep. 2019;9(1):1–16.

41. Berg S, Kutra D, Kroeger T, Straehle CN, Kausler BX, Haubold C, et al. ilastik: interactive machine learning for (bio)image analysis. Nat Methods. 2019;16(12):1226–32.

42. Schindelin J, Arganda-Carreras I, Frise E, Kaynig V, Longair M, Pietzsch T, et al. Fiji: An open-source platform for biological-image analysis. Nature Methods. 2012;9(7):676–82.

43. Zaimi A, Wabartha M, Herman V, Antonsanti PL, Perone CS, Cohen-Adad J. AxonDeepSeg: Automatic axon and myelin segmentation from microscopy data using convolutional neural networks. Sci Rep. 2018;8(1):1–11.

44. Kreshuk A, Straehle CN, Sommer C, Koethe U, Cantoni M. Automated Detection and Segmentation of Synaptic Contacts in Nearly Isotropic Serial Electron Microscopy Images. PLoS One. 2011;6(10):24899. doi: 10.1371/journal.pone.0024899.

45. Kreshuk A, Zhang C. Machine Learning: Advanced Image Segmentation Using ilastik. In: Rebollo, E., Bosch, M. (eds) Computer Optimized Microscopy. Methods in Molecular Biology. Humana, New York, NY; 2019. pp 449–463. doi: 10.1007/978-1-4939-9686-5_21.

46. Vergara HM, Pape C, Meechan KI, Zinchenko V, Genoud C, Wanner AA, et al. Whole-body integration of gene expression and single-cell morphology. Cell. 2021;184(18):4819-37.e22. doi: 10.1016/j.cell.2021.07.017.

47. Pachitariu M, Stringer C. Cellpose 2.0: how to train your own model. Nat Methods. 2022;19:1634–41.

48. Boyd A, Zhang H, Williams A. Insufficient OPC migration into demyelinated lesions is a cause of poor remyelination in MS and mouse models. Acta Neuropathol. 2013;125(6):841–59.

49. Landini G. Advanced shape analysis with ImageJ. In: Proceedings of the Second ImageJ User and Developer Conference. Luxembourg; Nov 6–7, 2008. p. 116–21. ISBN 2-919941-06-2.

50. Caicedo JC, Roth J, Goodman A, Becker T, Karhohs KW, Broisin M, et al. Evaluation of Deep Learning Strategies for Nucleus Segmentation in Fluorescence Images. Cytometry Part A. 2019; 95A:952–65. doi: 10.1002/cyto.a.23863.

51. Landini G, Galton A, Randell D, Fouad S. Novel applications of discrete mereotopology to mathematical morphology. Signal Process Image Commun. 2019;76:109–17.

